# Discovery of *Theta* Ribozymes in Gut Phages–Implications for tRNA and Alternative Genetic Coding

**DOI:** 10.1101/2023.06.13.544163

**Authors:** Kasimir Kienbeck, Lukas Malfertheiner, Susann Zelger-Paulus, Silke Johannsen, Christian von Mering, Roland K.O. Sigel

**Affiliations:** Department of Chemistry, University of Zurich; Zurich, CH-8057, Switzerland; Department of Molecular Life Sciences and Swiss Institute of Bioinformatics, University of Zurich; Zurich, CH-8057, Switzerland

**Author notes:** These authors contributed equally to this work (Kasimir Kienbeck, Lukas Malfertheiner).

## Abstract

Ribozymes, relics of the “RNA world”, are essential across all domains of life. Nonetheless, the functions and genomic contexts of recently discovered small ribozymes, such as minimal hepatitis delta virus (HDV)-like ribozymes, remain elusive. Using bioinformatic analyses, we identified a novel subfamily of minimal HDV-like ribozymes, coined *theta* ribozymes. Hundreds of unique examples were found adjacent to viral tRNAs within *Caudoviricetes* bacteriophages of the mammalian gut virome. *In vitro* experiments confirm site-specific self-scission activity, suggesting their involvement in processing tRNA 3’-trailers.

Intriguingly, a significant fraction of *theta* ribozymes is associated with viral suppressor tRNAs, potentially regulating the late-stage assembly of recoded bacteriophages. These findings advance the understanding of RNA-based mechanisms underlying the intricate interplay between the bacterial and viral parts of the mammalian gut microbiome.

**One-Sentence Summary:** Newly unveiled *theta* ribozymes associate with suppressor tRNAs of alternatively coded gut phages: a potential lytic switch.

## Main Text

Ribozymes are ubiquitous and participate in essential biological processes in all domains of life, including peptidyl transferase activity (*1*) and the transesterification steps required for tRNA maturation (*2*) as well as eukaryotic mRNA splicing (*3*). Small ribozymes (<200 nucleotides; nt) are restricted to self-cleavage and/or -ligation but are remarkably diverse in sequence, structure, and biological functions (*4*–*7*). A well-studied example is the family of hepatitis delta virus (HDV)-like ribozymes (delta-like ribozymes, drzs), which have a highly conserved, nested double-pseudoknotted structure but considerable variability in primary sequence (Fig. 1A) (*8*– *10*). While biological functions of specific drz examples are known (*11*–*16*), the majority, especially the minimal variants, are less understood. Minimal drzs, which were first identified in metagenomic samples, lack the P4 domain (Fig. 1A) (*17*) and their origin (eukaryotic, bacterial, or viral) and biological functions remain to be determined.

**Fig. 1.**
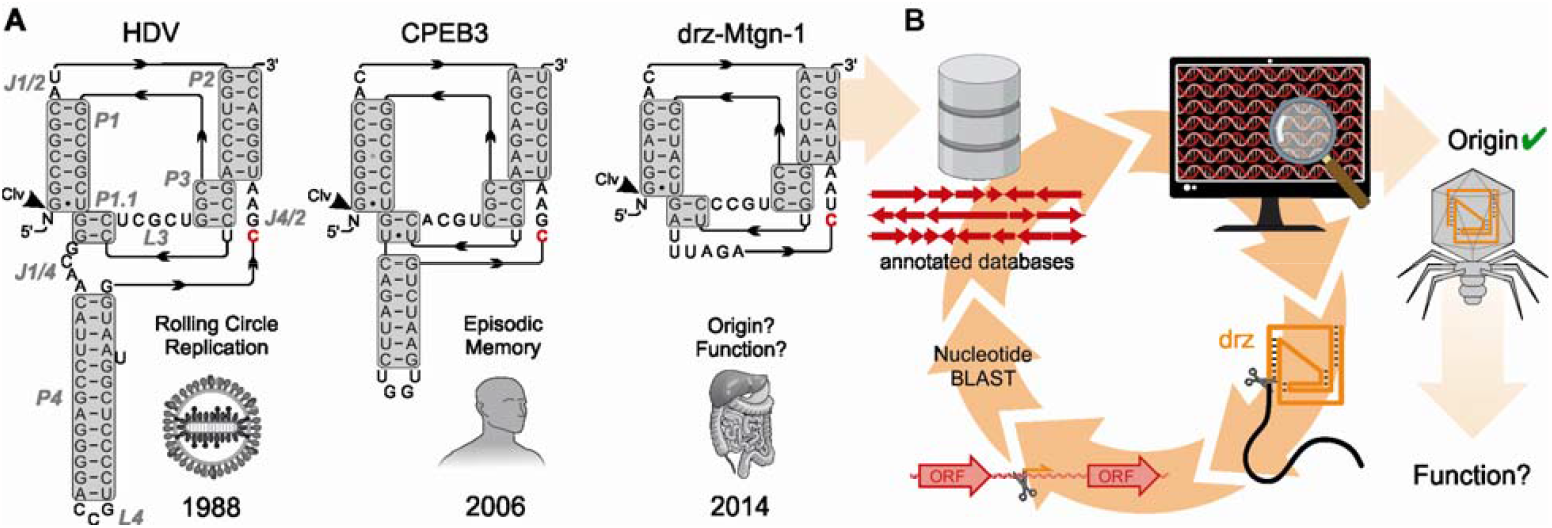
HDV-like ribozymes (drzs) and their association with bacteriophages. (**A**) Secondary structure of representative drzs: HDV (*11*), the human CPEB3 ribozyme (*54*), and the metagenomic drz “drz-Mtgn-1” with unknown origin and function (*17*). Helical domains are highlighted in gray, domain labels in gray italics, catalytic cytosine residues are marked in red. (**B**) Workflow of the initial nucleotide-based ribozyme homology search: BLAST analysis recovered drz-Mtgn-1 from annotated bacteriophage databases, subsequent BLAST analyses of nearby open reading frames (ORF) revealed novel ribozymes (fig. S1). Colors: viral DNA and proteins: red; drz: orange.

Herein, we report the discovery and *in vitro* validation of minimal drzs within *Caudoviricetes* bacteriophage genomes of the mammalian gut often associated with viral tRNAs and designate them as *theta* ribozymes (Θrzs). The gut virome is mainly composed of bacteriophages (>90%) and is increasingly linked to human health and disease (*18*–*21*), leading to multiple sequence database additions in recent years. Sequence analyses of these databases have revealed that some bacteriophages are recoded: they use genetic codes in which a certain stop-codon is reassigned to a standard amino acid (*22*). One example, where the amber stop-codon (UAG) is recoded to glutamine (code 15), was recently experimentally verified (*23*). Viral suppressor tRNAs (tRNA^Sup^) are central players in the translation of alternative genetic codes, a tool which may be used by recoded phages to initiate host lysis. However, the precise mechanism of the lytic-lysogenic switch in bacteriophages is not yet fully characterized.

In this study, we propose that tRNA-associated Θrzs are involved in viral tRNA maturation and regulate the code switch in a subset of recoded bacteriophages, potentially even triggering phage lysis. Our findings shed light on the poorly understood lytic-lysogenic switch in bacteriophages and provide new insights into the intriguing world of drzs and their biological significance.

### The viral origin of metagenomic minimal drzs

The minimal drz “drz-Mtgn-1” was identified in a metagenomic sample (*17*), but its origin and biological function remain unclear (Fig. 1A). We were intrigued by this knowledge gap and the drz’s unique behavior with divalent metal ions, and conducted a nucleotide sequence-based search of publicly available sequence databases (*24*). This search ultimately led, among others, to the assignment of drz-Mtgn-1 to several double-stranded DNA (dsDNA) bacteriophages (*Caudoviricetes*, Fig. 1B).

Due to the conservation of drz secondary structure rather than primary sequence, we changed our approach to motif-based RNArobo (*25*) searches using a minimal motif by Riccitelli, *et al*. (*17*). Initial searches resulted in over 60 novel minimal drz sequences in bacteriophage genomes (*26, 27*) and three conclusions: (i) minimal drz sequences were exclusively detected in bacteriophage genomes assembled from human gut metagenomic data, (ii) the hits showed associations with nearby open reading frames (ORF; mostly of proteins with unknown functions), potentially providing insights into their biological functions (fig. S1), and (iii) minimal drzs are more widespread than previously thought, with dozens of hits discovered in an initial search of two databases compared to a few hits from a full-scale search conducted in 2014 (*17*).

### Identification of tRNA-associated Θrzs in bacteriophages

A subset of hits within our initial categorization captured our attention, specifically, minimal drzs adjacent to phage tRNA genes. The position of the ribozyme cleavage site (G1; Fig 1A) at the 3’-end of the tRNA suggests a novel biological function in tRNA maturation. We therefore focused on these examples and refined our search motif accordingly. We chose eight recently annotated viral databases from diverse environments (table S1) and cross-referenced all subsequent hits with tRNA motif searches in the same databases. To increase motif specificity, we incorporated false-positive motifs as internal controls, considering previous findings that drzs are inactivated by a cytosine-to-uracil mutation (CΔU) at the active site (*28*), with no observed rescue mutation at this position (*29*). Each search was conducted with four different descriptor files, one active ribozyme motif with the catalytic cytosine residue intact (first position in the J4/2 junction) and three false-positive motifs containing substitutions at this residue (CΔA, CΔG, and CΔU, respectively; Fig. 2A). An initial search yielded less than 50% of the total hits with the active motif, indicating a high false positive rate (Fig. 2B (i)). We iteratively improved the motif by shortening the L4 loop and J1/2 junction (Fig. 2B (i), (ii)) and applying nucleotide identity constraints based on preliminary consensus sequences (>97% conservation; Fig. 2B (iii)). Finally, we introduced one additional degree of freedom at the last position of the J4/2 junction in line with the HDV-like structural motif (*9*). The latter reduced the false positive rate to nearly zero while increasing the number of detected tRNA-associated ribozymes (Fig. 2B (iv)). Sequences obtained from this refined motif are referred to as *theta* ribozymes (Θrz) due to their frequent associations with tRNAs.

**Fig. 2.**
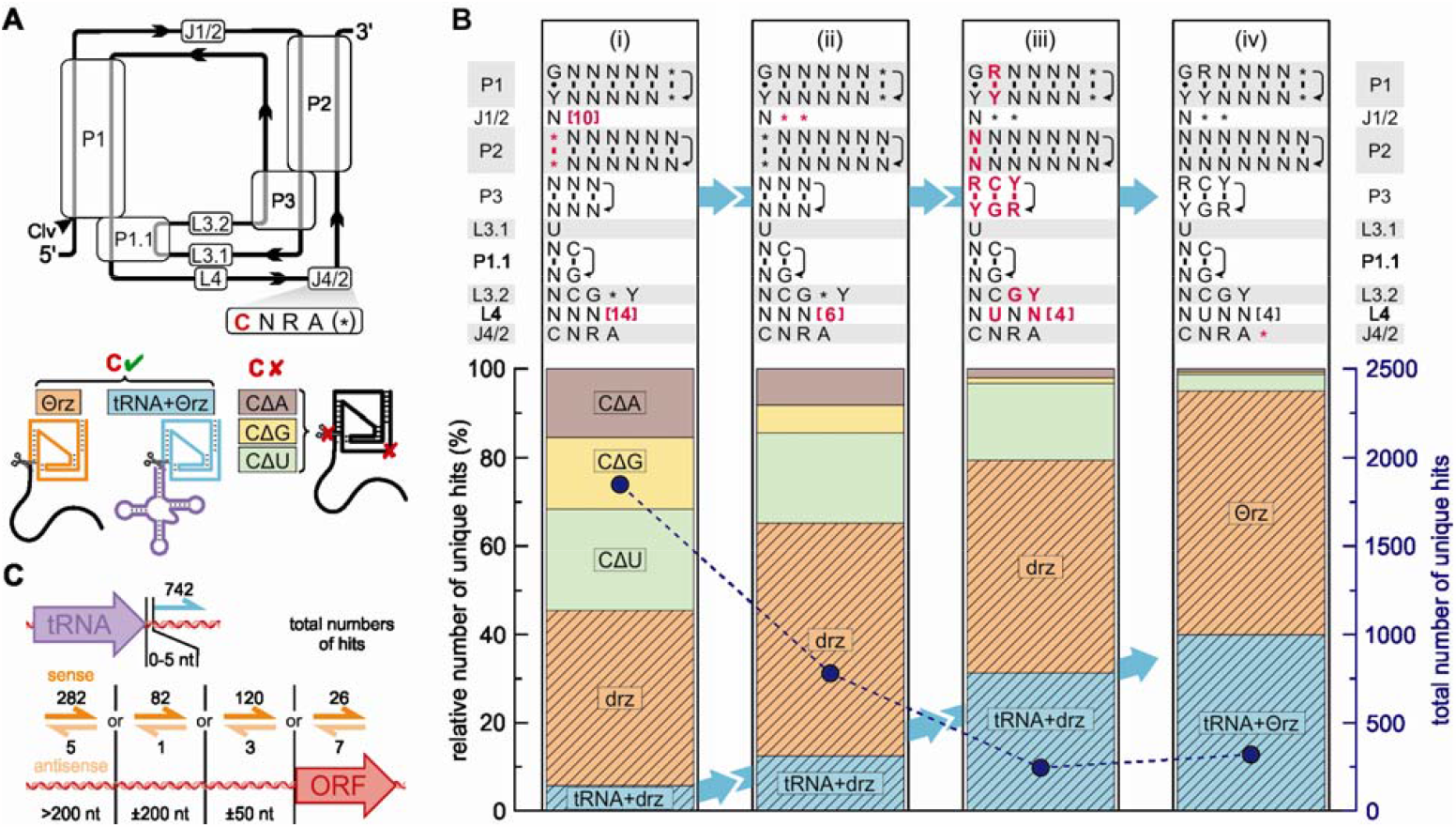
Optimization of the *theta* ribozyme (Θrz) motif using annotated bacteriophage databases. (**A**) Secondary structure of minimal drzs. The catalytic cytosine (red) in the J4/2 junction was substituted with A (CΔA: brown), G (CΔG: yellow), and U (CΔU: green), resulting in inactive motifs (false positives). True positives were subdivided into isolated ribozymes (orange) and tRNA-associated ribozymes (turquoise). (**B**) Changes to the search motif (top) in four steps (i) – (iv) and hits from annotated bacteriophage databases (bottom). Top: Domains denominated as in (A), helical strand directionality is indicated (black arrows). Modifications to the previous motif are indicated in bold red. (i) shows changes to the motif by Riccitelli *et al*. (*17*). IUPAC nucleotide denominations. * = any or no nucleotide. [X] = between zero and X nucleotides. Bottom: Relative number of unique hits with the motif above (left y-axis). Putatively active ribozyme hits are striped. Total hit numbers shown in marine blue (right y-axis; points and dashed line). Ribozyme sequences obtained by the final motif (iv) are denominated as Θrzs. (**C**) Top: Total tRNA-associated Θrzs (turquoise half-arrows) adjacent to viral tRNAs (purple). Bottom: Total number of isolated Θrz hits (orange half-arrows and black numbers) in sense (dark orange) or antisense (light orange) to the closest ORF (red arrow).

Using the optimized motif, we identified 302 unique Θrz sequences in the above-mentioned databases, with 126 classified as tRNA-associated Θrzs, where the ribozyme’s cleavage site (G1) was within ±5 nt of the 3’-end of a tRNA. Our analysis revealed 152 distinct Θrz-adjacent tRNA sequences, resulting in 185 unique tRNA/Θrz combinations. Many Θrzs are present in multiple viral genomes, totaling 1281 occurrences, with 742 (58%) classified as tRNA-associated Θrzs (Fig. 2C, top). 568 of these tRNA-associated Θrzs (77%) are directly adjacent to the respective tRNA (±1 nt).

Over 80% of Θrzs were found in human or animal gut bacteriophages, with no or only a few hits in other environments (table S1). Our analysis included bacterial and eukaryotic genomes (human, mouse, and protists), revealing only 12 Θrz hits in bacterial genomes (*30*), none of which were associated with a tRNA. Consistent with our initial manual searches, nearly 99% of Θrz sequences were confined to bacteriophages belonging to the *Caudoviricetes* order, the predominant family of dsDNA viruses in the human gut virome (fig. S2) (*31*).

The non-tRNA-associated Θrz hits (n=526) were categorized based on their closest ORF (Fig. 2C, bottom). The majority (n=493) reside in non-coding regions, most being located more than 200 nt from the nearest annotated ORF. Over 96% of these Θrz hits shared the same directionality (sense) as the closest up- or downstream gene. Only 33 examples were located partially or entirely within an ORF (intragenic). Considering the anticipated self-cleavage of identified ribozymes in an HDV-like manner, we expect them to be located outside of coding regions, indicating that the non-tRNA associated Θrzs likely serve unknown biological functions. However, tRNA-associated Θrzs constitute the majority of our hit pool and demonstrate the clearest indication of a biological function, prompting us to focus on this sub-family for *in vitro* validation.

### tRNA-associated Θrzs are active *in vitro*

The internal transesterification mechanism of drzs relies on an essential cytosine in the J4/2 linker with a perturbed p*K*_a_ and a coordinated Mg^2+^ ion, which positions the phosphate backbone for an in-line attack. This acid-base catalyzed reaction has been shown to be most efficient near neutral pH (*32*). To validate the HDV-like self-scission of tRNA-associated Θrzs, we selected and investigated four tRNA/Θrz pairs *in vitro* (fig. S3). We chose three pairs of higher prevalence in the initial search and the tRNA^Val^0025_Θ0046 (for naming see methods) pair because of its elongated J4/2 junction, which allowed us to experimentally verify the additional introduced degree of freedom in the final motif as true positives (Fig. 2B (iv)).

All selected examples exhibited Mg^2+^-dependent self-scission activity *in vitro* (Fig. 3A and B and fig. S4A). The apparent self-cleavage rate constant (*k*_obs_) showed a typical HDV-like sigmoidal behavior with increasing Mg^2+^ concentration. Among the constructs, the tRNA/Θrz pair tRNA^Val^0025_Θ0046 exhibited the highest *k*_obs_ at pH 7.0 (> 10 min^−1^ at 10 mM Mg^2+^; Fig. 3B and D), while the other constructs showed significantly slower *k*_obs_ at the same Mg^2+^ concentration (10 mM, Fig. 3D and table S2).

**Fig. 3.**
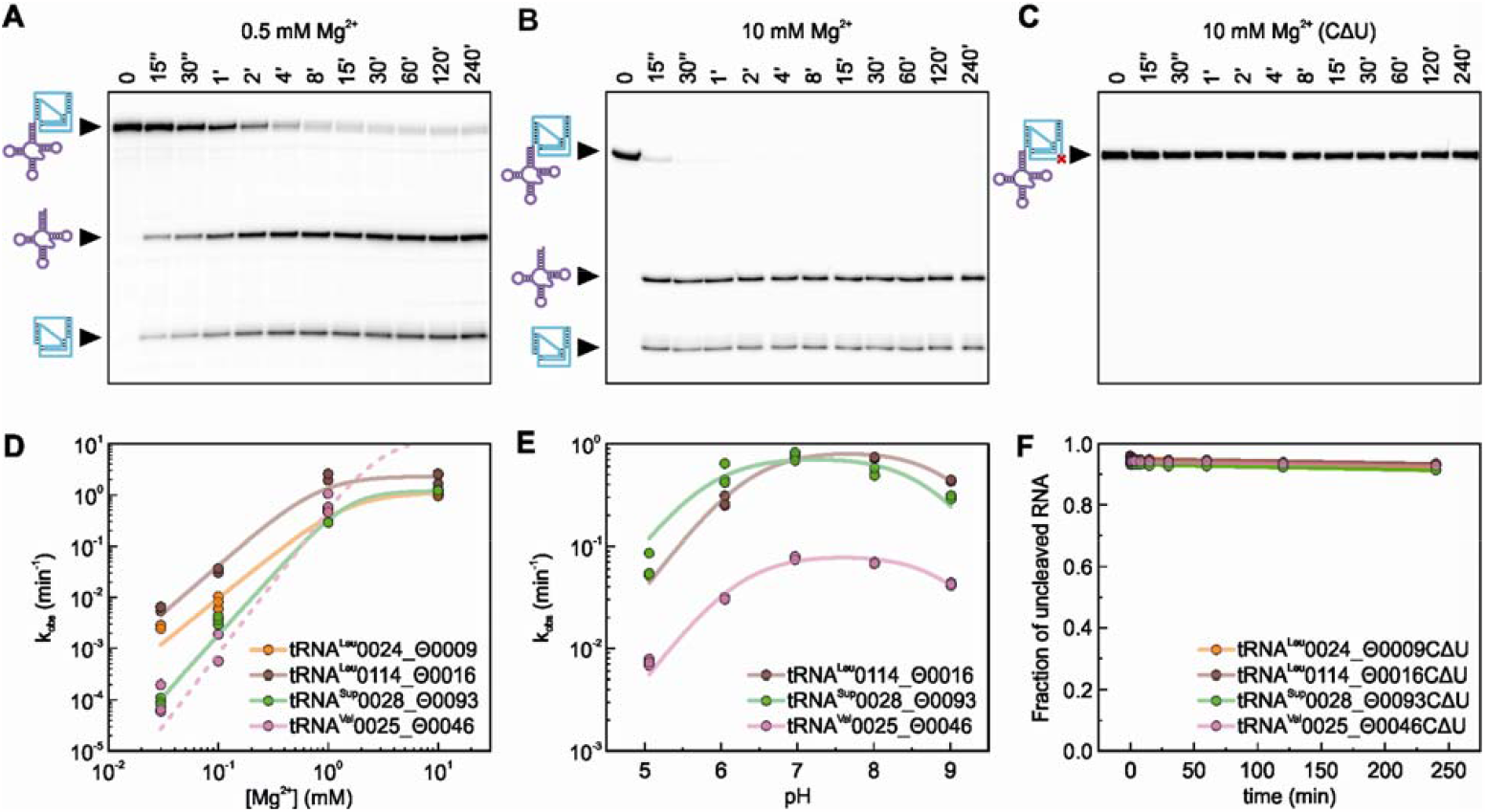
*In vitro* confirmation of selected tRNA/Θrz pairs. (**A**) Representative PAGE analysis of self-cleavage assays with tRNA^Val^0025_Θ0046 RNA at pH 7.0 and 37 °C induced with 0.5 mM Mg^2+^. Timepoints are indicated above the respective lanes (‘‘ = seconds; ‘ = minutes). Band identities: Precursor RNA = combined cartoon (146 nt); tRNA = purple (90 nt); Θrz = turquoise (56 nt). (**B**) Representative gel of tRNA^Val^0025_Θ0046 RNA at pH 7.5 and 37 °C induced with 10 mM Mg^2+^. (**C**) Representative gel of tRNA^Val^0025_Θ0046CΔU RNA at pH 7.5 and 37 °C induced with 10 mM Mg^2+^. (**D**) Calculated apparent kinetic rate constants (*k*_obs_) at varying Mg^2+^ concentrations of the four examples tRNA^Leu^0024_Θ0009, tRNA^Leu^0114_Θ0016, tRNA^Sup^0028_Θ0093, and tRNA^Val^0025_Θ0046. The *k*_obs_-Mg^2+^ dependency for tRNA^Val^0025_Θ0046 at 10 mM Mg^2+^ can only be estimated due to too fast cleavage (dashed line). (**E**) Calculated *k*_obs_ at varying pH values of the three examples tRNA^Leu^0114_Θ0016, tRNA^Sup^0028_Θ0093, and tRNA^Val^0025_Θ0046. (**F**) Relative fraction of precursor RNA of the four inactivated examples (tRNA^Leu^0024_Θ0009CΔU, tRNA^Leu^0114_Θ0016CΔU, tRNA^Sup^0028_Θ0093CΔU, and tRNA^Val^0025_Θ0046CΔU) at 37°C induced with 10 mM Mg^2+^.

Titrations of three tRNA/Θrz pairs across a pH range of 5 to 9 revealed maximal *k*_obs_ at physiological pH, consistent with previously identified minimal drzs (Fig. 3E and fig. S4B) (*17*). Two p*K*_a_ values were inferred for each construct: p*K*_a1_ ≈ 9.0 probably corresponding to a hydrated Mg^2+^ ion and p*K*_a2_ ≈ 6.0 related to the catalytic cytosine residue (table S3).

Furthermore, we confirmed the inactivation of all tRNA/Θrz pairs upon mutating the catalytic cytosine to uracil (CΔU), in line with known HDV-like behavior (*29*). This observation further validates our approach for identifying false-positives in the bioinformatic search (Fig. 3C and F). In summary, these *in vitro* self-cleavage assays do not show any unexpected behavior but confirm the HDV-like nature of Θrzs.

### Enrichment of tRNA^Sup^-associated Θrzs reveals recoded bacteriophages

To gain insights into the prevalence of Θrzs outside of annotated databases, we conducted an extensive search on raw reads from 469’049 publicly available metagenomic datasets using our improved search motif. This search yielded additional 104’264 hits (9’344 unique Θrz sequences), of which 12’158 (11.7%) were tRNA-associated, resulting in 1’698 unique Θrzs adjacent to 5’721 unique tRNAs. Consistent with previous findings from bacteriophage databases, the human gastrointestinal tract emerged as the primary environment for Θrzs (fig. S5).

To give a comprehensive overview, tables sorted by Θrz frequency in descending order are provided in data S1 and S2. These data include combined Θrz sequences and their adjacent tRNA sequences from annotated and metagenomic samples with additional information (data S1, n=13’257), as well as non-tRNA-associated Θrzs (data S2, n=7’704). Remarkably, we found Θrzs associated with predicted tRNAs of all amino acids (Fig. 4A). Among them, 157 unique Θrzs were associated with tRNAs of more than one amino acid isotype (excluding undetermined types; Undet), with Θ0013 displaying the greatest diversity (13 different amino acid isotypes).

**Fig. 4.**
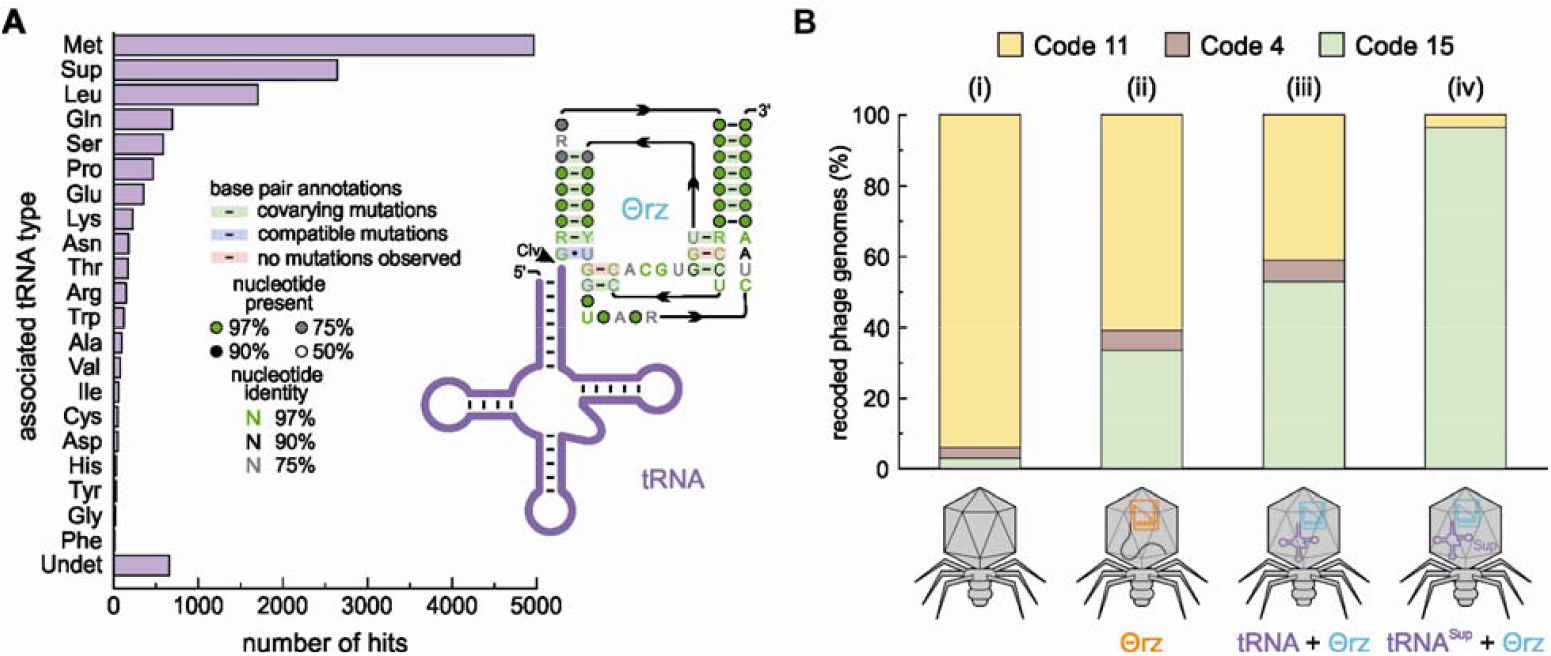
Categorization of tRNA-associated Θrz hits and probability of recoding in phage genomes. (**A**) Total number of Θrz hits (n=13’257) from combined annotated and metagenomic datasets sorted by the type of associated tRNA located ± 5 nt from the Θrz cleavage site. Undet = undetermined tRNA type. A consensus sequence of all unique tRNA-associated Θrz hits (n=1’753) was visualized using R2R (*33*). Nucleotide denominations according to IUPAC nomenclature. (**B**) Fraction of annotated bacteriophage genomes (%) predicted to use code 15 (UAG = Gln; green), code 4 (UGA = Trp; brown), or the standard bacterial genetic code 11 (yellow). (i) The overall expected recoding ratio of phages, (ii) phages containing Θrzs (orange), (iii) phages containing tRNA-associated Θrzs (turquoise) linked to a tRNA of any type (purple), and (iv) phages containing Θrzs linked to tRNA^Sup^.

Alignment and analysis of all unique tRNA-associated Θrz sequences using R2R (*33*) resulted in a consensus motif featuring several conserved nucleotides not originally defined in the descriptor file. These hits were further categorized based on the associated tRNA type, and over 70% of all hits are associated with either tRNA^Met^, tRNA^Sup^, or tRNA^Leu^ (Fig. 4A). Our interest was piqued especially by the large proportion of Θrzs (20%) associated with tRNA^Sup^, of which 99.7% contain an anticodon for the amber stop codon (UAG). These results suggest stop-codon reassignments, since tRNA^Sup^ are essential elements of recoded phages and are required for successful phage lysis in cases where the host employs a different genetic code (*22*). These findings may provide answers to a fundamental open question: Why do bacteriophages invest resources in carrying tRNAs instead of utilizing host-provided tRNAs?

Building upon recent findings by Borges *et al*. (*22*), who reported stop-codon reassignment in ∼2-6% of human and animal gut phages (Fig. 4B (i)), we investigated the genetic codes of the analyzed bacteriophages. Our predictions comprise the predominant recodings code 15 (UAG reassigned to Gln), code 4 (UGA reassigned to Trp), and the standard bacterial code (code 11). Genomes with a reassigned stop-codon show gene fragmentation when genes are predicted in standard code. Therefore, we classified a phage genome as recoded if the alternative coding density exceeded a 5-10% threshold (depending on the genome size) compared to the coding density in the standard code. Remarkably, we found that over one-third (33.7%) of genomes containing Θrz sequences likely utilize code 15, and 5.6% use code 4 (Fig. 4B (ii)). When we narrowed down our analysis to genomes containing tRNA-associated Θrzs, these proportions increased to over half (53.0%) and 6.0%, respectively (Figure 4B, (iii)). Notably, when analyzing only phage genomes containing tRNA^Sup^-associated Θrzs, a remarkable 96.4% are likely recoded to code 15 (no hits of code 4 were observed; Fig. 4B, (iv)). Therefore, we expect the tRNA^Sup^ of these phages to show a glutamine isotype. However, computational isotype predictions are based solely on bacterial tRNAs (*34*) and require further experimental validation to draw definitive conclusions. Nonetheless, our subsequent investigation of the 180 tRNA^Sup^ examples yielded 69.4% of tRNA^Sup^ with a predicted glutamine isotype, and the remainder isotypes for Trp (19.4%; rarely reported code 32), Ile (10.0%; unreported recoding), and Tyr (1.1%; code 29). Phage genome analysis and tRNA^Sup^ predictions both point to code 15 recoding, strengthening the hypothesis that Θrzs play an essential role in the code switch of recoded phages.

## Discussion and outlook

An iterative improvement of an existing minimal drz motif (*17*) using bacteriophage genomes resulted in a reliable motif with a low false positive rate and high specificity for tRNA-associated Θrzs (Fig. 2). By restricting the optimization process to unique Θrz sequences we also increased stringency, resulting in the identification of over 100’000 Θrzs in metagenomic and annotated databases. Although a short length of most raw reads makes the detection of both a Θrz and tRNA on the same read highly unlikely, we still detect a notable proportion (∼13%) of tRNA-associated examples. Thus, the true number of tRNA-associated Θrzs is likely much larger than reported.

Our biochemical analyses confirmed HDV-like self-scission *in vitro* for all four selected ribozymes. Moreover, their efficient self-scission rates are comparable to or even faster than those of previously reported metagenomic examples (e.g. *k*_obs_(drz-Mtgn-3) = 1.69 ± 0.03 min^−1^ and *k*_obs_(drz-Mtgn-4) = 0.0022 ± 0.0001 min^−1^) (*17*). These properties make them potential candidates for bioengineering and biomedical applications, such as aptazymes: self-cleaving ribozymes combined with aptamers to control gene expressions (*35*–*38*).

Due to the frequent association of Θrzs with tRNA encoding sequences, we postulate a function in tRNA 3’-trailer processing, which has not been reported to date and is currently limited to *Caudoviricetes* bacteriophages in mammalian gut microbiomes. While the generation of the mature 5’-end of tRNAs is well-understood and involves a single ribonucleoprotein enzyme (RNase P) present in all domains of life (*39*), the 3’-processing of tRNAs is less understood. In *Escherichia coli*, the cleavage of tRNA 3’-trailers involves a complex, multi-step mechanism involving various endo-(RNases E and III) and exonucleases (RNases II, BN, D, PH, PNPase, and T) (*40*). We postulate that bacteriophages containing Θrzs do not need to rely on host RNases, the production of which may be suppressed. By associating Θrzs in *cis* with tRNAs instead of encoding the various nucleases itself, the phage significantly reduces the genomic space required to produce mature tRNA 3’-ends, despite the need for a ribozyme at each tRNA. We hypothesize that this adaptation contributes to phage fitness and explains the prevalence of tRNA-associated Θrzs observed in this order.

The regulation of tRNAs is crucial not only for protein biosynthesis, but also for host-manipulation during viral infections: recent evidence has revealed that viral tRNAs can substitute cellular tRNAs and support viral infection (*41*). Combined with the requirement of viral tRNAs to sustain translation while the host machinery degrades (*42*), these factors may explain the prevalence of tRNAs in viral genomes. With respect to tRNA^Sup^, which are essential for recoded phages, we show a clear positive correlation between tRNA^Sup^-associated Θrzs and code 15 phages. The remarkably high abundance and efficiency of Θrzs provide additional support and a new key element in the mechanism recently proposed by Borges *et al*.: tRNA^Sup^-associated Θrzs might regulate or even initiate the lytic cycle in specific bacteriophages (Fig. 5). Their study highlighted the overrepresentation of stop codons in phage structural and lysis genes (*22*), suggesting a pivotal role for stop-codon reassignment in the timing and mechanism of the lysis trigger. Importantly, mistiming can be detrimental, as premature lysis serves as a host defense mechanism. In such a scenario, the host initiates lysis before phage particles are fully assembled within the cell, severely compromising phage efficiency (*43*–*45*).

**Fig. 5.**
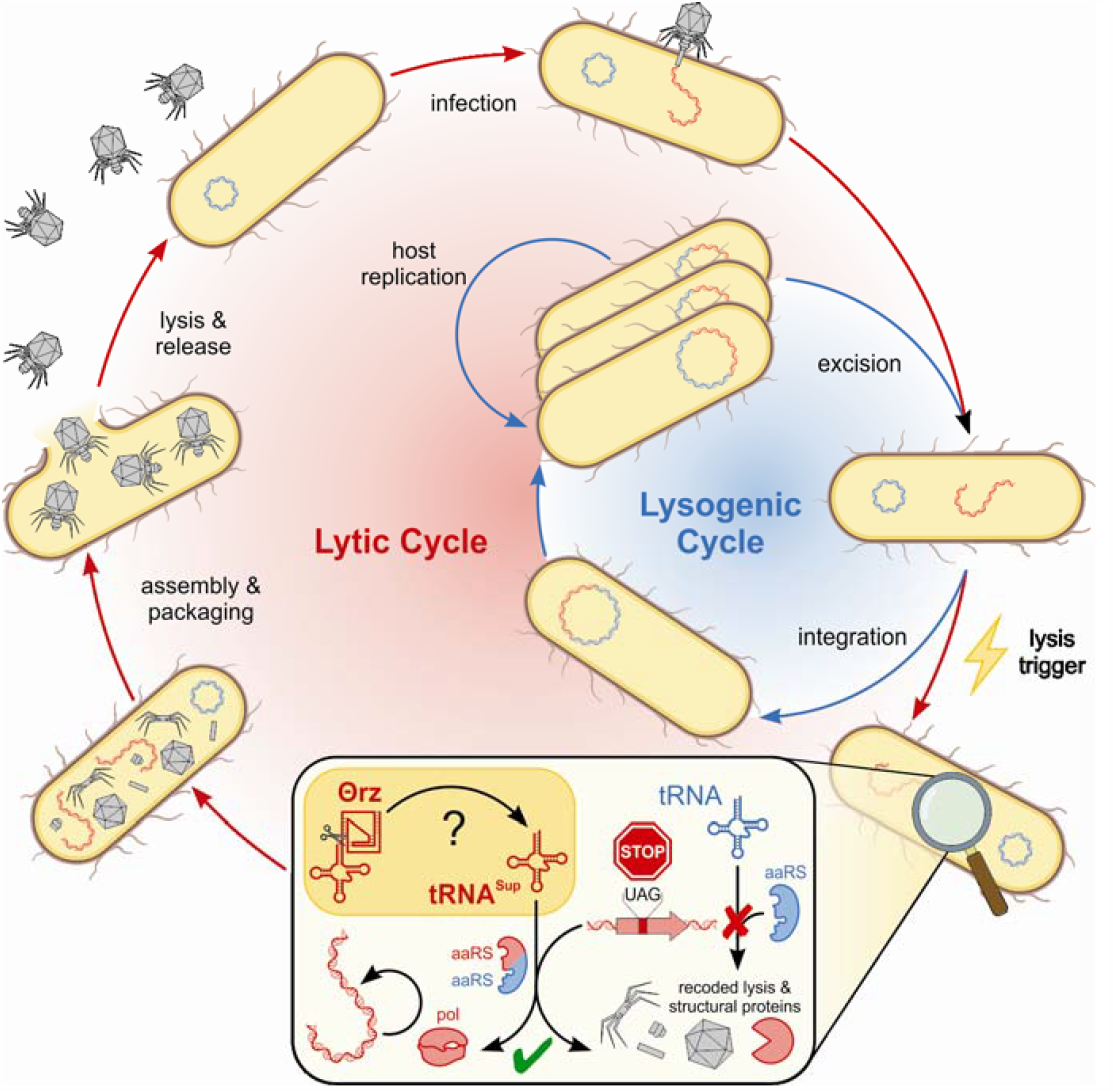
Putative phage infection cycle involving tRNA^Sup^-associated Θrzs. A gut bacteria is infected by a recoded *Caudoviricetes* bacteriophage, which can initiate the lysogenic or lytic cycle. In the lysogenic cycle, the phage genome is integrated into the host genome and host-encoded machinery replicates the host cell including the integrated phage. A not fully understood mechanism triggers the lytic cycle (lightning symbol). Phage encoded tRNA^Sup^, possibly regulated by associated Θrzs, enable the translation of recoded lysis and structural proteins (enlarged box). An attempt to produce these proteins using host-encoded tRNAs and aminoacyl-tRNA-synthetases (aaRS) would lead to gene fragmentation (in-frame amber stop-codons). In the last phase of the lytic cycle, phage particles self-assemble, the host cell is lysed, and phage particles are released, leading to a new infection cycle. Polymerase: pol. Colors: Host DNA/proteins: blue; phage DNA/RNA/proteins: red.

The exact activation and regulation of Θrz self-scission as well as the release of the associated tRNA *in vivo* is still unclear and subject to future studies. An alternative hypothesis not involving the necessity of direct Θrz regulation could be that the ribozyme persists in an “always on” state and self-regulates the translation of late phase lytic genes in a concentration-based manner. This would involve highly efficient co-transcriptional tRNA 3’-trailer scission by the Θrz, drastically increasing the concentration of tRNA^Sup^ within the host cell. Once a critical concentration is surpassed, the viral tRNA^Sup^ overwhelms host-encoded termination factors and leads to the code switch from code 11 to 15, enabling late-phase viral gene translation and subsequent bacterial host cell lysis. A similar regulation is observed for ribonucleotide reductase, which reduces ribonucleotides to deoxyribonucleotides. This enzyme is activated or deactivated based on the concentration and therefore preferential binding of ATP or dATP to the allosteric activity site, respectively (*46, 47*). The observed association of Θrz sequences with multiple tRNA isotypes further strengthens concentration-based self-regulation. Θrzs may not only initiate a code switch but also optimize the codon usage for phage-encoded genes, further supporting their translation rather than host-encoded genes.

By providing the sequences of all identified Θrzs and their associated tRNAs, we offer a valuable resource with thousands of examples for future studies. Furthermore, we report the discovery of tRNA^Sup^ sequences with a predicted Ile-isotype, representing an unobserved recoding event.

Overall, our combined *in silico* and *in vitro* results emphasize the importance of ribozymes in biological processes and introduce a new subfamily of small tRNA processing ribozymes, which enable extensive tRNA-mediated host manipulations within the mammalian gut microbiome.

## Supporting information

Supplementary Materials including Materials and Methods, Figs. S1 to S6, and Tables S1 to S3

## Acknowledgements

We are grateful to Andrej Luptak, the Sigel and von Mering lab group members for fruitful discussions in the course of this study.

## Author contributions

Conceptualization: KK, LM, SZP, RKO

Methodology: KK, LM, SZP, SJ, CvM, RKOS

Investigation: KK, LM

Software: LM, KK

Data curation: LM

Visualization: KK

Funding acquisition: SZP, RKOS, CvM

Project administration: SZP, SJ, RKOS, CvM

Supervision: SZP, SJ, CvM, RKOS

Writing – original draft: KK, LM

Writing – review & editing: KK, LM, SZP, SJ, RKOS, CvM

## Data and materials availability

All data are available in the main text or the supplementary materials. The viral databases from open sources were obtained from the following links:

https://zenodo.org/record/4776317 (*27*)

http://ftp.ebi.ac.uk/pub/databases/metagenomics/genome_sets/gut_phage_database/ (*48*)

http://www.virusite.org/index.php?nav=download (*49*)

https://zenodo.org/record/6410225 (*22*)

https://portal.nersc.gov/MGV (*50*)

https://github.com/RChGO/OVD/ (*51*)

https://datacommons.cyverse.org/browse/iplant/home/shared/iVirus/GOV2.0 (*52*)

https://genome.jgi.doe.gov/portal/IMG_VR/IMG_VR.home.html (*53*)

## Supplementary Materials

Materials and Methods

Figs. S1 to S6

Tables S1 to S3

Data S1 to S3

## References and Notes

1. P. Nissen, J. Hansen, N. Ban, P. B. Moore, T. A. Steitz, The Structural Basis of Ribosome Activity in Peptide Bond Synthesis. Science. 289, 920–930 (2000).

2. C. Guerrier-Takada, K. Gardiner, T. Marsh, N. Pace, S. Altman, The RNA Moiety of Ribonuclease P is the Catalytic Subunit of the Enzyme. Cell. 35, 849–857 (1983).

3. M. E. Wilkinson, C. Charenton, K. Nagai, RNA Splicing by the Spliceosome. Annu. Rev. Biochem. 89, 359–388 (2020).

4. J. A. Doudna, J. R. Lorsch, Ribozyme catalysis: not different, just worse. Nat. Struct. Mol. Biol. 12, 395–402 (2005).

5. M. J. Fedor, J. R. Williamson, The catalytic diversity of RNAs. Nat. Rev. Mol. Cell Biol. 6, 399–412 (2005).

6. P. C. Bevilacqua, R. Yajima, Nucleobase catalysis in ribozyme mechanism. Curr. Opin. Chem. Biol. 10, 455–464 (2006).

7. W. G. Scott, Ribozymes. Curr. Opin. Struct. Biol. 17, 280–286 (2007).

8. C.-H. T. Webb, N. J. Riccitelli, D. J. Ruminski, A. Lupták, Widespread Occurrence of Self-Cleaving Ribozymes. Science. 326, 953–953 (2009).

9. C.-H. T. Webb, A. Lupták, HDV-like self-cleaving ribozymes. RNA Biol. 8, 719–727 (2011).

10. N. J. Riccitelli, A. Lupták, Computational discovery of folded RNA domains in genomes and in vitro selected libraries. Methods San Diego Calif. 52, 133–140 (2010).

11. L. Sharmeen, M. Y. P. Kuo, G. Dinter-Gottlieb, J. Taylor, Antigenomic RNA of Human Hepatitis Delta Virus Can Undergo Self-Cleavage. J. Virol. 62, 2674–2679 (1988).

12. C. Vogler, K. Spalek, A. Aerni, P. Demougin, A. Müller, K.-D. Huynh, A. Papassotiropoul, D. J.-F. de Quervain, CPEB3 is associated with human episodic memory. Front. Behav. Neurosci. 3, 1–5 (2009).

13. D. J. Ruminski, C.-H. T. Webb, N. J. Riccitelli, A. Lupták, Processing and Translation Initiation of Non-long Terminal Repeat Retrotransposons by Hepatitis Delta Virus (HDV)-like Self-cleaving Ribozymes. J. Biol. Chem. 286, 41286–41295 (2011).

14. D. G. Eickbush, T. H. Eickbush, R2 Retrotransposons Encode a Self-Cleaving Ribozyme for Processing from an rRNA Cotranscript. Mol. Cell. Biol. 30, 3142–3150 (2010).

15. F. J. Sánchez-Luque, M. C. López, F. Macias, C. Alonso, M. C. Thomas, Identification of an hepatitis delta virus-like ribozyme at the mRNA 5′-end of the L1Tc retrotransposon from Trypanosoma cruzi. Nucleic Acids Res. 39, 8065–8077 (2011).

16. J. L. Jakubczak, W. D. Burke, T. H. Eickbush, Retrotransposable elements R1 and R2 interrupt the rRNA genes of most insects. Proc. Natl. Acad. Sci. U. S. A. 88, 3295–3299 (1991).

17. N. J. Riccitelli, E. Delwart, A. Lupták, Identification of Minimal HDV-Like Ribozymes with Unique Divalent Metal Ion Dependence in the Human Microbiome. Biochemistry. 53, 1616–1626 (2014).

18. Z. Cao, N. Sugimura, E. Burgermeister, M. P. Ebert, T. Zuo, P. Lan, The gut virome: A new microbiome component in health and disease. eBioMedicine. 81, 104113 (2022).

19. A. G. Clooney, T. D. S. Sutton, A. N. Shkoporov, R. K. Holohan, K. M. Daly, O. O’Regan, F. J. Ryan, L. A. Draper, S. E. Plevy, R. P. Ross, C. Hill, Whole-Virome Analysis Sheds Light on Viral Dark Matter in Inflammatory Bowel Disease. Cell Host Microbe. 26, 764–778 (2019).

20. Y. Duan, R. Young, B. Schnabl, Bacteriophages and their potential for treatment of gastrointestinal diseases. Nat. Rev. Gastroenterol. Hepatol. 19, 135–144 (2022).

21. G. Nakatsu, H. Zhou, W. K. K. Wu, S. H. Wong, O. O. Coker, Z. Dai, X. Li, C.-H. Szeto, N. Sugimura, T. Y.-T. Lam, A. C.-S. Yu, X. Wang, Z. Chen, M. C.-S. Wong, S. C. Ng, M. T. V. Chan, P. K. S. Chan, F. K. L. Chan, J. J.-Y. Sung, J. Yu, Alterations in Enteric Virome Are Associated With Colorectal Cancer and Survival Outcomes. Gastroenterology. 155, 529–541 (2018).

22. A. L. Borges, Y. C. Lou, R. Sachdeva, B. Al-Shayeb, P. I. Penev, A. L. Jaffe, S. Lei, J. M. Santini, J. F. Banfield, Widespread stop-codon recoding in bacteriophages may regulate translation of lytic genes. Nat. Microbiol. 7, 918–927 (2022).

23. S. L. Peters, A. L. Borges, R. J. Giannone, M. J. Morowitz, J. F. Banfield, R. L. Hettich, Experimental validation that human microbiome phages use alternative genetic coding. Nat. Commun. 13, 5710 (2022).

24. E. W. Sayers, E. E. Bolton, J. R. Brister, K. Canese, J. Chan, D. C. Comeau, C. M. Farrell, M. Feldgarden, A. M. Fine, K. Funk, E. Hatcher, S. Kannan, C. Kelly, S. Kim, W. Klimke, M. J. Landrum, S. Lathrop, Z. Lu, T. L. Madden, A. Malheiro, A. Marchler-Bauer, T. D. Murphy, L. Phan, S. Pujar, S. H. Rangwala, V. A. Schneider, T. Tse, J. Wang, J. Ye, B. W. Trawick, K. D. Pruitt, S. T. Sherry, Database resources of the National Center for Biotechnology Information in 2023. Nucleic Acids Res. 51, D29–D38 (2023).

25. L. Rampášek, R. M. Jimenez, A. Lupták, T. Vinař, B. Brejová, RNA motif search with data-driven element ordering. BMC Bioinformatics. 17, 216 (2016).

26. S. Nishijima, N. Nagata, Y. Kiguchi, Y. Kojima, T. Miyoshi-Akiyama, M. Kimura, M. Ohsugi, K. Ueki, S. Oka, M. Mizokami, T. Itoi, T. Kawai, N. Uemura, M. Hattori, Extensive gut virome variation and its associations with host and environmental factors in a population-level cohort. Nat. Commun. 13, 5252 (2022).

27. M. J. Tisza, C. B. Buck, A catalog of tens of thousands of viruses from human metagenomes reveals hidden associations with chronic diseases. Proc. Natl. Acad. Sci. 118, e2023202118 (2021).

28. A. Ke, K. Zhou, F. Ding, J. H. D. Cate, J. A. Doudna, A conformational switch controls hepatitis delta virus ribozyme catalysis. Nature. 429, 201–205 (2004).

29. J. M. Roberts, J. D. Beck, T. B. Pollock, D. P. Bendixsen, E. J. Hayden, RNA sequence to structure analysis from comprehensive pairwise mutagenesis of multiple self-cleaving ribozymes. eLife. 12, e80360 (2023).

30. A. Fullam, I. Letunic, T. S. B. Schmidt, Q. R. Ducarmon, N. Karcher, S. Khedkar, M. Kuhn, M. Larralde, O. M. Maistrenko, L. Malfertheiner, A. Milanese, J. F. M. Rodrigues, C. Sanchis-López, C. Schudoma, D. Szklarczyk, S. Sunagawa, G. Zeller, J. Huerta-Cepas, C. von Mering, P. Bork, D. R. Mende, proGenomes3: approaching one million accurately and consistently annotated high-quality prokaryotic genomes. Nucleic Acids Res. 51, D760–D766 (2023).

31. G. Liang, F. D. Bushman, The human virome: assembly, composition and host interactions. Nat. Rev. Microbiol. 19, 514–527 (2021).

32. S. Nakano, D. M. Chadalavada, P. C. Bevilacqua, General Acid-Base Catalysis in the Mechanism of a Hepatitis Delta Virus Ribozyme. Science. 287, 1493–1497 (2000).

33. Z. Weinberg, R. R. Breaker, R2R--software to speed the depiction of aesthetic consensus RNA secondary structures. BMC Bioinformatics. 12, 3 (2011).

34. P. P. Chan, B. Y. Lin, A. J. Mak, T. M. Lowe, tRNAscan-SE 2.0: improved detection and functional classification of transfer RNA genes. Nucleic Acids Res. 49, 9077–9096 (2021).

35. J. Stifel, M. Spöring, J. S. Hartig, Expanding the toolbox of synthetic riboswitches with guanine-dependent aptazymes. Synth. Biol. Oxf. Engl. 4, ysy022 (2019).

36. G. Zhong, H. Wang, C. C. Bailey, G. Gao, M. Farzan, Rational design of aptazyme riboswitches for efficient control of gene expression in mammalian cells. eLife. 5, e18858 (2016).

37. H. Peng, B. Latifi, S. Müller, A. Lupták, I. A. Chen, Self-cleaving ribozymes: substrate specificity and synthetic biology applications. RSC Chem. Biol. 2, 1370–1383 (2021).

38. S. Kobori, K. Takahashi, Y. Yokobayashi, Deep Sequencing Analysis of Aptazyme Variants Based on a Pistol Ribozyme. ACS Synth. Biol. 6, 1283–1288 (2017).

39. J. C. Ellis, J. W. Brown, The RNase P family. RNA Biol. 6, 362–369 (2009).

40. S. Schiffer, S. Rösch, A. Marchfelder, Assigning a function to a conserved group of proteins: the tRNA 3′-processing enzymes. EMBO J. 21, 2769–2777 (2002).

41. A. Nyerges, S. Vinke, R. Flynn, S. V. Owen, E. A. Rand, B. Budnik, E. Keen, K. Narasimhan, J. A. Marchand, M. Baas-Thomas, M. Liu, K. Chen, A. Chiappino-Pepe, F. Hu, M. Baym, G. M. Church, A swapped genetic code prevents viral infections and gene transfer. Nature. 615, 720–727 (2023).

42. J. Y. Yang, W. Fang, F. Miranda-Sanchez, J. M. Brown, K. M. Kauffman, C. M. Acevero, D. P. Bartel, M. F. Polz, L. Kelly, Degradation of host translational machinery drives tRNA acquisition in viruses. Cell Syst. 12, 771–779 (2021).

43. E. Durmaz, T. R. Klaenhammer, Abortive phage resistance mechanism AbiZ speeds the lysis clock to cause premature lysis of phage-infected Lactococcus lactis. J. Bacteriol. 189, 1417–1425 (2007).

44. S. G. Hays, K. D. Seed, Dominant Vibrio cholerae phage exhibits lysis inhibition sensitive to disruption by a defensive phage satellite. eLife. 9, e53200 (2020).

45. R. Johnson-Boaz, C.-Y. Chang, R. Young, A dominant mutation in the bacteriophage lambda S gene causes premature lysis and an absolute defective plating phenotype. Mol. Microbiol. 13, 495–504 (1994).

46. A. Hofer, M. Crona, D. T. Logan, B.-M. Sjöberg, DNA building blocks: keeping control of manufacture. Crit. Rev. Biochem. Mol. Biol. 47, 50–63 (2012).

47. B. L. Greene, G. Kang, C. Cui, M. Bennati, D. G. Nocera, C. L. Drennan, J. Stubbe, Ribonucleotide Reductases (RNRs): Structure, chemistry, and metabolism suggest new therapeutic targets. Annu. Rev. Biochem. 89, 45–75 (2020).

48. L. F. Camarillo-Guerrero, A. Almeida, G. Rangel-Pineros, R. D. Finn, T. D. Lawley, Massive expansion of human gut bacteriophage diversity. Cell. 184, 1098–1109 (2021).

49. M. Stano, G. Beke, L. Klucar, viruSITE-integrated database for viral genomics. Database J. Biol. Databases Curation. 2016, baw162 (2016).

50. S. Nayfach, D. Páez-Espino, L. Call, S. J. Low, H. Sberro, N. N. Ivanova, A. D. Proal, M. Fischbach, A. S. Bhatt, P. Hugenholtz, N. C. Kyrpides, Metagenomic compendium of 189,680 DNA viruses from the human gut microbiome. Nat. Microbiol. 6, 960–970 (2021).

51. S. Li, R. Guo, Y. Zhang, P. Li, F. Chen, X. Wang, J. Li, Z. Jie, Q. Lv, H. Jin, G. Wang, Q. Yan, A catalog of 48,425 nonredundant viruses from oral metagenomes expands the horizon of the human oral virome. iScience. 25, 104418 (2022).

52. A. C. Gregory, A. A. Zayed, N. Conceição-Neto, B. Temperton, B. Bolduc, A. Alberti, M. Ardyna, K. Arkhipova, M. Carmichael, C. Cruaud, C. Dimier, G. Domínguez-Huerta, J. Ferland, S. Kandels, Y. Liu, C. Marec, S. Pesant, M. Picheral, S. Pisarev, J. Poulain, J.-É. Tremblay, D. Vik, Tara Oceans Coordinators, M. Babin, C. Bowler, A. I. Culley, C. de Vargas, B. E. Dutilh, D. Iudicone, L. Karp-Boss, S. Roux, S. Sunagawa, P. Wincker, M. B. Sullivan, Marine DNA Viral Macro- and Microdiversity from Pole to Pole. Cell. 177, 1109–1123 (2019).

53. S. Roux, D. Páez-Espino, I.-M. A. Chen, K. Palaniappan, A. Ratner, K. Chu, T. B. K. Reddy, S. Nayfach, F. Schulz, L. Call, R. Y. Neches, T. Woyke, N. N. Ivanova, E. A. Eloe-Fadrosh, N. C. Kyrpides, IMG/VR v3: an integrated ecological and evolutionary framework for interrogating genomes of uncultivated viruses. Nucleic Acids Res. 49, D764–D775 (2021).

54. K. Salehi-Ashtiani, A. Lupták, A. Litovchick, J. W. Szostak, A Genomewide Search for Ribozymes Reveals an HDV-Like Sequence in the Human CPEB3 Gene. Science. 313, 1788–1792 (2006).

55. N. A. O’Leary, M. W. Wright, J. R. Brister, S. Ciufo, D. Haddad, R. McVeigh, B. Rajput, B. Robbertse, B. Smith-White, D. Ako-Adjei, A. Astashyn, A. Badretdin, Y. Bao, O. Blinkova, V. Brover, V. Chetvernin, J. Choi, E. Cox, O. Ermolaeva, C. M. Farrell, T. Goldfarb, T. Gupta, D. Haft, E. Hatcher, W. Hlavina, V. S. Joardar, V. K. Kodali, W. Li, D. Maglott, P. Masterson, K. M. McGarvey, M. R. Murphy, K. O’Neill, S. Pujar, S. H. Rangwala, D. Rausch, L. D. Riddick, C. Schoch, A. Shkeda, S. S. Storz, H. Sun, F. Thibaud-Nissen, I. Tolstoy, R. E. Tully, A. R. Vatsan, C. Wallin, D. Webb, W. Wu, M. J. Landrum, A. Kimchi, T. Tatusova, M. DiCuccio, P. Kitts, T. D. Murphy, K. D. Pruitt, Reference sequence (RefSeq) database at NCBI: current status, taxonomic expansion, and functional annotation. Nucleic Acids Res. 44, D733–D745 (2016).

56. D. Hyatt, G.-L. Chen, P. F. Locascio, M. L. Land, F. W. Larimer, L. J. Hauser, Prodigal: prokaryotic gene recognition and translation initiation site identification. BMC Bioinformatics. 11, 119 (2010).

57. A. P. Camargo, S. Roux, F. Schulz, M. Babinski, Y. Xu, B. Hu, P. S. G. Chain, S. Nayfach, N. C. Kyrpides, You can move, but you can’t hide: identification of mobile genetic elements with geNomad (2023), (available at https://www.biorxiv.org/content/10.1101/2023.03.05.531206v1).

58. A. Mueller, J.-C. Fillion-Robin, R. Boidol RCCG, P. Nechifor, 田丰收, J. Berdat, A. Keleg, P. Hunt yoonsubKim, B. Petrushev, C. Minshall, E. Guan, Y. Dai, J. Medina, M. Corvellec, R. Rampin, H. Xu, K. M. Langner, K. Bhamidipati, C. Gieringer, I. Ozsvald, Beppe, freeIsa cjmay D. Roy, D. Logvinenko IgorAPM, T. Sato, V. Rho, WordCloud 1.5.0 (2023), (available at https://zenodo.org/record/1322068).

