## Supplementary Materials including Materials and Methods, Figs. S1 to S6, and Tables S1 to S3 for "Discovery of *Theta* Ribozymes in Gut Phages–Implications for tRNA and Alternative Genetic Coding"

**Affiliations:**

**The file includes:**

Materials and Methods
Figs. S1 to S6
Tables S1 to S3

**Other Supplementary Materials for this manuscript include the following:**

Data S1 to S3

### Materials and Methods

Initial BLAST searches

Using the sequence "GGTAGCACACCTATGCGTTCCCGTCGCGCTACTGATTTAGACTAAATAGGT" as a query (drz-Mtgn-1 (*17*)), initial manual homology searches (BLASTn) were conducted against the nucleotide collection databases from NCBI (*24*). Several hits with 100% identity were observed, nearby ORFs were screened manually and submitted to further manual homology searches. Flanking regions of ORFs with an e-value cutoff of $10^{-10}$ were investigated by hand initially. Subsequently, the corresponding databases were downloaded and submitted to motif searches using the software RNArobo (*25*) (see below for detailed description). This procedure was repeated several times, resulting in dozens of hits, among them a first hint of tRNA associations (fig. S1).

Motif-based database searches and motif improvement

Eukaryotic genomes were chosen from NCBI RefSeq (*55*) with the following query title on June 10th, 2023:

'Search Eukaryota AND "complete genome"[filter] AND all[filter] NOT anomalous[filter].'

Additionally, human (GCA_000001405.28) and mouse (GCA_000001635.9) genomes were manually added to the genome collection. For Bacteria, the representative genomes from ProGenomes3 (*30*) (https://progenomes.embl.de/download.cgi) were downloaded. The viral databases were obtained on June 10th, 2022, under the links specified in Data availability (*22*, *26*, *27*, *48*–*53*). All files were merged for subsequent analysis. Due to some redundancies in the Roux, 2021 database (*53*) (contained three of our other databases: Nayfach, 2021 (*50*); Nishijima, 2022 (*26*); Gregory, 2019 (*52*)), only hits that were unique were considered for downstream analysis, *i.e.*, ribozymes that were already detected in other databases were discarded from that source. The RNArobo software (*25*) v2.1.0 was used to search these databases and define the search motif containing the conserved sequence and structural elements of minimal drzs. This motif was based on a minimal motif described by Riccitelli *et al.* (*17*), and the descriptor file was structured as shown in fig. S6.

The final descriptor file resulted from several iterations of searches on annotated phage genome databases (table S1), where the motif was manually adapted to result in a high percentage of Θrzs within the total drz hit pool. All databases were searched with RNArobo 2.1.0 and -c --nratio 0.1 parameters a total number of 16 times: 4 iterations with 4 different motifs (once the active motif and thrice the false-positive motifs). All motif iterations used are depicted in Fig. 2.

tRNAs were detected using tRNAscan-SE v2.0.9 (*34*) in general mode (-G). Due to computational time constraints, only contigs containing at least one drz hit were screened for tRNAs. Custom code in Python v.3.7.6 was used to combine all outputs and detect Θrzs that are located adjacent to tRNAs. A Θrz is considered tRNA-associated if a tRNA 3'-end was detected within ± 5 nucleotides of its cleavage site.

To obtain the predicted isotype tRNAscan-SE 2.0.9 was run in bacterial mode (-B -s), and the output was processed in a script in Python v.3.7.6. All code will be supplied upon publication of the final manuscript.

Metagenomic sequence search

All raw reads from sequence runs available in the MicrobeAtlas project (microbeatlas.org) marked as whole genome sequences were used in the subsequent analysis. Raw sequences were downloaded, and quality filtered as described under https://microbeatlas.org/index.html?action=help. All 469'049 sample runs matching the criteria were analyzed with both RNArobo and tRNA Scan-SE as described above. The sequence read archive (SRA) run IDs of the analyzed samples are stored in data S3.

The final table containing all tRNA-associated Θrz sequences was constructed using pandas v1.0.3. Both tables (metagenomic as well as database hits) were combined, and the ribozymes were sorted and named according to their occurrence. For example, $tRNA^{Val}$0025_Θ0046 RNA would be the combination of the $25^{th}$ most common tRNA and the $46^{th}$ most common tRNA-associated Θrz. Additionally, the source (sample run ID from SRA, or Database source and identifier) is provided in data S1. All unique isolated Θrzs were additionally collected, and subsequently named according to their number of occurences and uploaded in data S2.

<u>Coding density analysis</u>

Prodigal v2.6.3 (*56*) was used to infer coding sequences based on the three codes 11, 15, and 4 (-g11, -g15, -g4, respectively). The script get_CD.py from Borges *et al.* (*22*) (https://github.com/borgesadair1/AC_phage_analysis/releases/tag/v1.0.0) was adapted to calculate coding density for all genomes in the combined database. The coding density of a bacteriophage genome had to be at least 5 or 10% (depending on contig size: <100 kb: 10%, >100 kbp: 5%) higher with the alternative code than standard code to be considered a recoded phage.

<u>Taxonomy</u>

The taxonomy of all phage genomes was determined using genomad v1.3.3 (*57*) with the "genomad end-to-end -cleanup" workflow. A taxonomy was considered true if at least 75% of the genes agreed in their taxonomic assignments. All contigs containing Θrzs were separately analyzed with the same parameters, and their relative abundances of taxonomic assignments were compared and plotted with OriginPro, Version 2022.

<u>Word clouds</u>

Samples in the MicrobeAtlas Project are annotated with one of four main environments (animal, aquatic, soil, or plant) and keywords extracted from the metadata of the SRA. These environmental assignments and keywords can be found in the file "samples.env.info" obtained from https://microbeatlas.org/index.html?action=download, on March $10^{th}$, 2023. Custom code in Python v.3.7.6 was used to extract the environment and all keywords of the respective sample of each detected tRNA-associated Θrz. The list of the obtained keywords was used to create a word cloud with WordCloud v1.5.0 (*58*) with a custom color map and the following parameters:

stopwords=stopwords, prefer_horizontal=1, min_font_size=10, max_font_size=150, relative_scaling=.4, width=1000, collocations=False,height=400, max_words=15, random_state=1, background_color="white".  Additionally, a "background" expectancy of keywords was generated by repeating the process with 10000 random metagenomic samples.

The environments of all detected tRNA-associated Θrz were extracted and combined, excluding samples annotated with "aquatic, waste water" since it can be contaminated with human stool samples, thus not accurately representing the aquatic habitat. The same was repeated for all possible samples to obtain a background expectancy.

<u>R2R alignments</u>

All unique Θrz sequences were transformed into a Stockholm 1.0 format file using a custom Python 3 script in Jupyterlab v3.5.0. R2R v1.0.6 (*33*) was used with the following parameters: First, the consensus file was created as follows (filenames in square brackets):

--GSC-weighted-consensus [input].sto [consensus_file].cons.sto 3 0.97 0.9 0.75 4 0.97 0.9 0.75 0.5 0.1

Then, the output was generated utilizing a meta file pointing to the consensus file:

--disable-usage-warning [meta_file].r2r_meta [output].pdf

The output .pdf file was edited with CorelDRAW X7 v17.1.0.572.

DNA template preparation

The plasmid backbone used for *in vitro* transcription was derived from pJD20, kindly provided by Dr Anna Marie Pyle. Double-stranded synthetic DNA containing an *EcoR*I restriction site, the T7 promoter sequence (TAATACGACTCACTATA), a transcription start site (GGGAGA) followed by the tRNA/Θrz pair, a *Pst*I restriction site, and a *Hind*III restriction site, was obtained from Azenta Life Sciences. The synthetic DNA was digested with *EcoR*I and *Hind*III and cloned into the plasmid backbone digested with the same restriction enzymes. To obtain the transcription template, the plasmid was either linearized with *Pst*I or the region of interest amplified by PCR using a forward primer (GAATTCTAATACGACTCACTATAGGGA) and individual reverse primers.

RNA *in vitro* transcription

$^{32}$P body-labeled RNA was prepared by *in vitro* transcription at 37°C for 4-5 hours in 50-200 µL reaction volumes containing 40 mM Tris-HCl pH 7.5, 40 mM DTT, 2 mM spermidine, 5 mM each ATP, GTP, and UTP, 0.5 mM CTP, 0.01% triton X-100, 10-100 nM template, 10 mM $MgCl_2$, 20-30 µM inhibitor oligo (Microsynth), 0.1-1 µCi/µL [α-$^{32}$P]-CTP (Perkin Elmer and Hartmann Analytic) and an appropriate amount of T7 RNA polymerase (purified in house). The inhibitor oligos were designed individually for each tRNA/Θrz pair (CLC Main Workbench v23.0.2) to span the self-cleavage site with a targeted melting temperature of 50°C to reduce co-transcriptional scission.

The reaction was quenched with an equal volume of loading buffer (0.16% bromophenol blue, 10 mM EDTA pH 8.0 in formamide) and loaded onto an 8% denaturing polyacrylamide gel (29:1; Fisher bioreagents). After electrophoresis, the gels were exposed to a phosphorimage screen (FUJI MS 50340272), visualized using a Typhoon FLA 9500 Scanner (control software v1.1) and the bands corresponding to the full-length RNA excised using a clean, sterile scalpel. The RNA was eluted from the crushed gel slice for 4-6 hours at 4°C in 5 volumes of crush & soak buffer (10 mM MOPS pH 6.0, 1 mM EDTA, and 250 mM NaCl). To precipitate the RNA, 1/10$^{th}$ of the volume of 3 M sodium acetate pH 5.2 and 3 volumes of ice-cold, absolute Ethanol were added, and the mixture was incubated at -20°C overnight. The pelleted RNA was washed with 70% ethanol and dissolved in 50 µL $ddH_2O$.

Self-scission kinetics

The kinetic reaction buffer (140 mM KCl, 10 mM NaCl, and 50 mM Tris-HCl pH 7.5) was pre-warmed at 37°C prior to the addition of purified $^{32}$P-labeled RNA to a final concentration of 0.5-1 nM after liquid scintillation counting (HIDEX 300SL; MikroWin v4.44). The reaction mixture was equilibrated at 37°C for 5 minutes, and self-scission was initiated by the addition of a 10-fold concentrated $MgCl_2$ stock solution to the desired concentration. 10 µL aliquots were taken at predetermined time points, quenched with equal volumes of loading buffer and loaded onto an 8% denaturing PAGE. After electrophoresis, the gel was dried (Whatman Biometra Maxidry D64), exposed onto a phosphorimage screen overnight and visualized using a Typhoon Scanner. Quantification of the bands was performed using ImageQuant TL v8.2.

In the pH titrations, the kinetic reaction buffer was replaced by two separate three-buffer systems depending on the pH range to guarantee constant ionic strength at all pH values. For pH 4.5-7.5, a buffer containing 25 mM MES (2-(*N*-morpholino)-ethanesulfonic acid), 25 mM acetic acid, 50 mM Tris (tris(hydroxymethyl)-aminomethane), 10 mM NaCl, and 140 mM KCl was used, whereas for pH 7.5-9.5, a buffer containing 50 mM MES, 25 mM Tris, 25 mM 2-amino-2-methyl-1-propanol, 10 mM NaCl, and 140 mM KCl was used.

Data fitting

The data fitting was performed using OriginPro, v2022. The band intensities of the cleaved tRNA bands were corrected for the number of cytosine residues and the relative intensities of $f_{tRNA}$ were calculated as follows (Eq. 1):

$$f_{tRNA} = \frac{I_{tRNA}}{I_{tRNA} + I_{substrate}} \quad (1)$$

$I_{tRNA}$ and $I_{substrate}$ correspond to the intensities of the tRNA and substrate (tRNA_Θrz RNA), respectively. The obtained values were fit to either an inverted mono- (Eq. 2) or biexponential decay function (Eq. 3):

$$f_{tRNA} = 1 - \left(A * e^{(-k_1 * t)} + C\right) \quad (2)$$

or

$$f_{tRNA} = 1 - \left(A * e^{(-k_1 * t)} + B * e^{(-k_2 * t)} + C\right) \quad (3)$$

*A*, *B* and *C* correspond to the relative fractions of the constructs performing self-scission at the rates $k_1$, $k_2$, or none, respectively. The reported cleavage rates ($k_{obs}$) correspond to $k_1$ from monoexponential and to the faster cleavage rate from biexponential fits.

The cleavage rate-$Mg^{2+}$-relationships were fit to the following Hill-equation (Eq. 4) assuming a single binding event resulting in self-scission with rate $k_1$:

$$k_{obs} = \frac{k_{\max}}{1 + \left(\frac{K_d}{\left[Mg^{2+}\right]}\right)^n} \quad (4)$$

This allows for the Hill coefficient *n*.

The cleavage rate-pH-relationships were fit to the following equation (Eq. 5) (*17*) assuming two titratable groups, namely a hydrated $Mg^{2+}$ ion ($pK_{a1}$) and the catalytic cytosine ($pK_{a2}$):

$$k_{obs} = \frac{k_{\max}}{1 + 10^{pH - pK_{a1}} + 10^{pK_{a2} - pH} + 10^{pKa2 - pK_{a1}}} \quad (5)$$

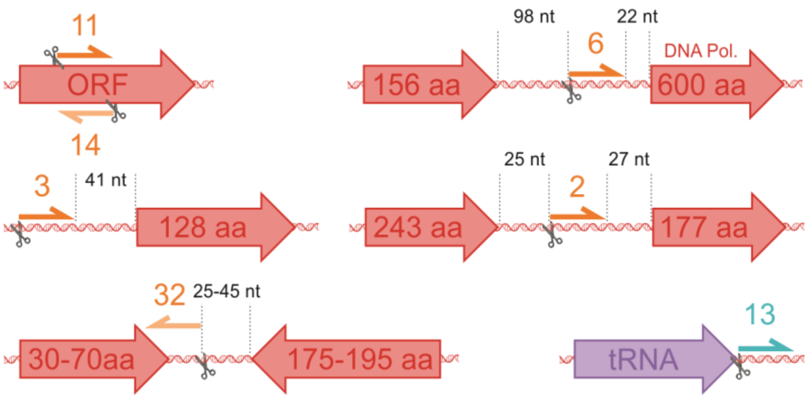


**Fig. S1. Initial classification of minimal drzs.** Initial classification of minimal drzs (orange half-arrows and numbers; orientation from 5' to 3') with respect to nearby annotated viral open reading frames (ORF; red arrows). DNA polymerase encoding ORF: DNA Pol.; Self-scission site of drzs indicated by scissors; distance to ORFs indicated in number of nucleotides (nt; black); ORF lengths indicated in number of amino acids (aa); viral tRNA encoding sequence: purple; tRNA-associated drzs: turquoise; drzs in the same orientation as ORFs (sense) depicted in dark colors; drzs in the opposite orientation as ORFs (antisense) depicted in light colors.

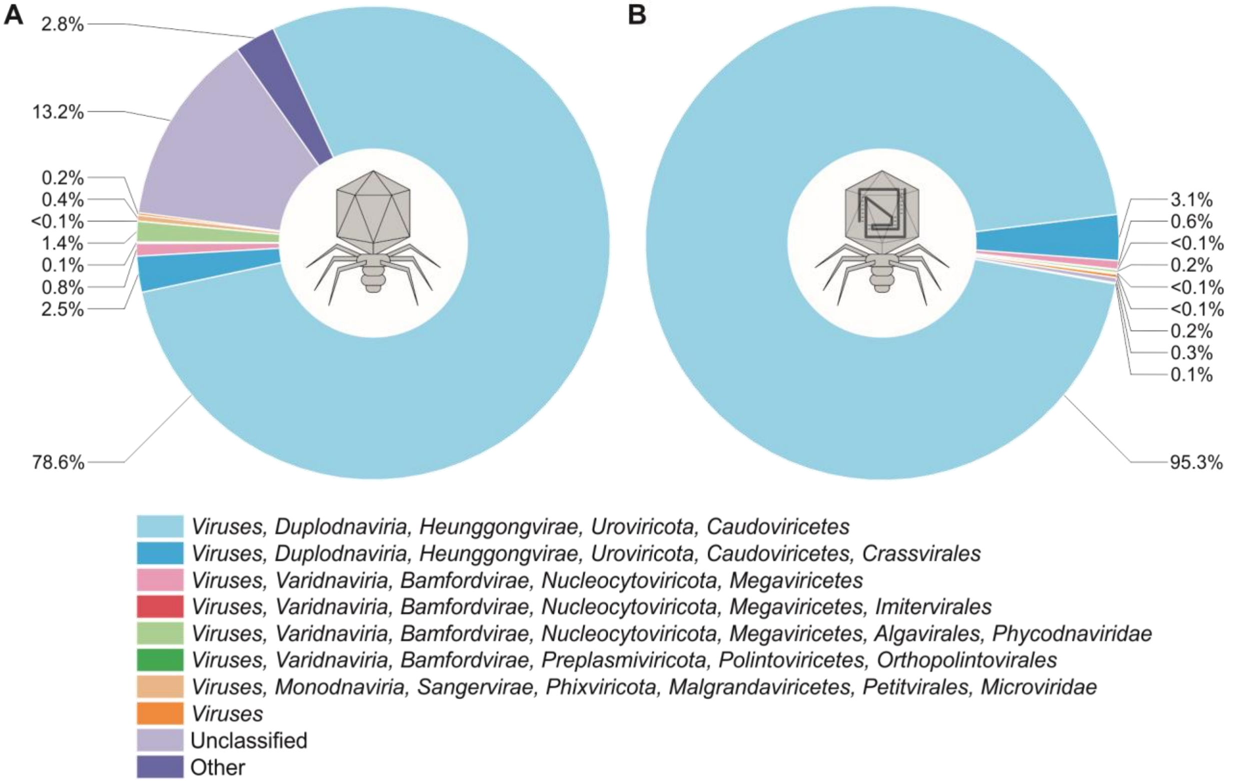


**Fig. S2. Phage taxonomy.** (**A**) Taxonomy of all viruses that were searched. (**B**) Taxonomy of viruses containing a Θrz. Almost all viruses (>97%) with an identified Θrz belong to the *Caudoviricetes*.

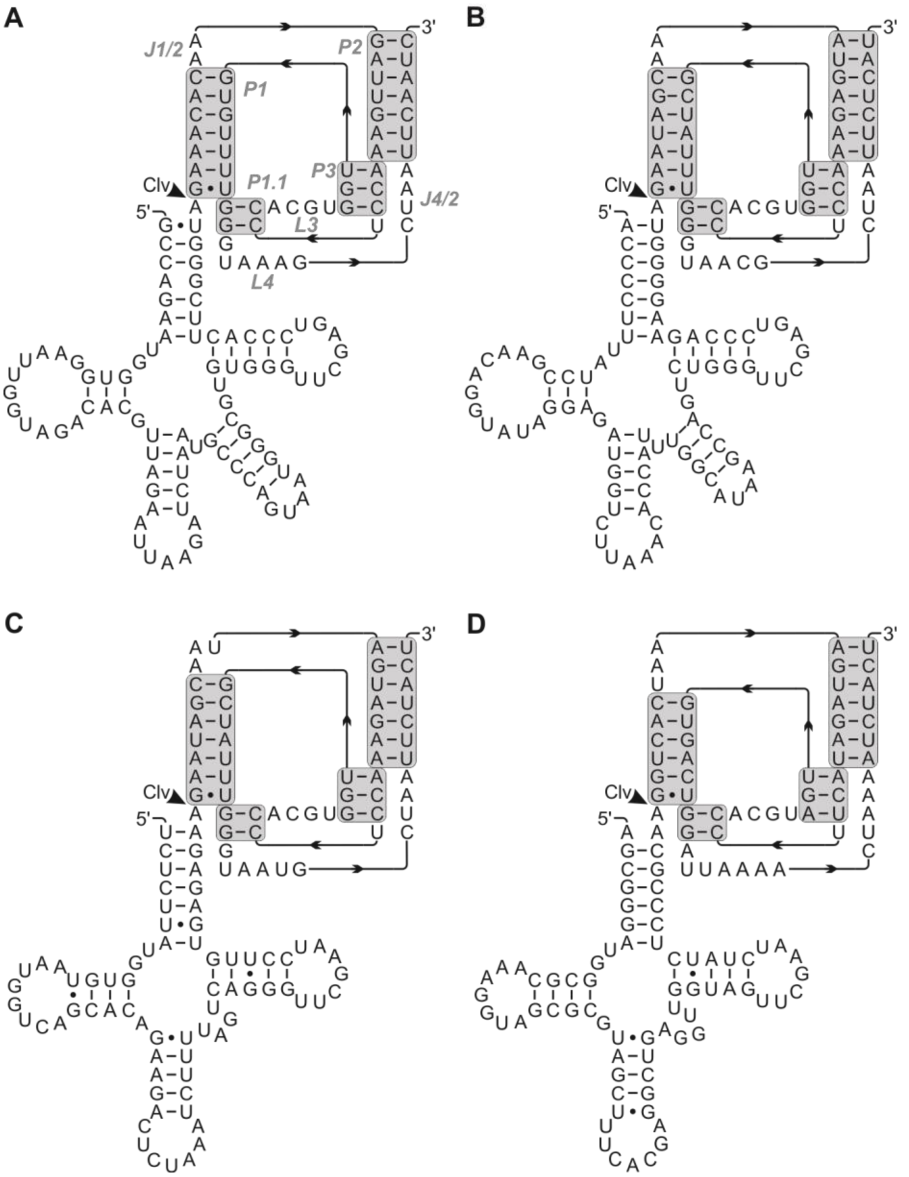


**Fig. S3. Secondary structures of analyzed tRNA/Θrz pairs.** (**A**) Proposed secondary structure of $\text{tRNA}^{\text{Leu}}$0024_Θ0009. Helical domains of the Θrz are highlighted in gray. Ribozyme domain labels in gray italics. (**B**) Proposed secondary structure of $\text{tRNA}^{\text{Leu}}$0114_Θ0016, labeling as in (A). (**C**) Proposed secondary structure of $\text{tRNA}^{\text{Sup}}$0028_Θ0093, labeling as in (A). (**D**) Proposed secondary structure of $\text{tRNA}^{\text{Val}}$0025_Θ0046, labeling as in (A). Cleavage site (Clv) is indicated by an arrowhead.

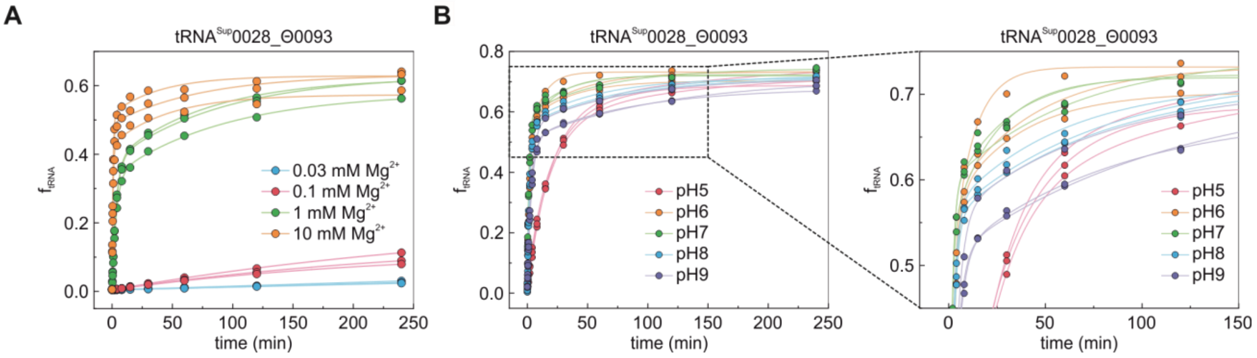


**Fig. S4. Representative fits of tRNA$^{Sup}$0028_Θ0093 self-cleavage.** (**A**) Magnesium titration of $f_{tRNA}$ calculated with Eq. 1. Curve was fitted using Eqs. 2 and 3. (**B**) pH titration of $f_{tRNA}$ calculated with Eq. 1 and enlargement for clarity. Curve was fitted using Eqs. 2 and 3.

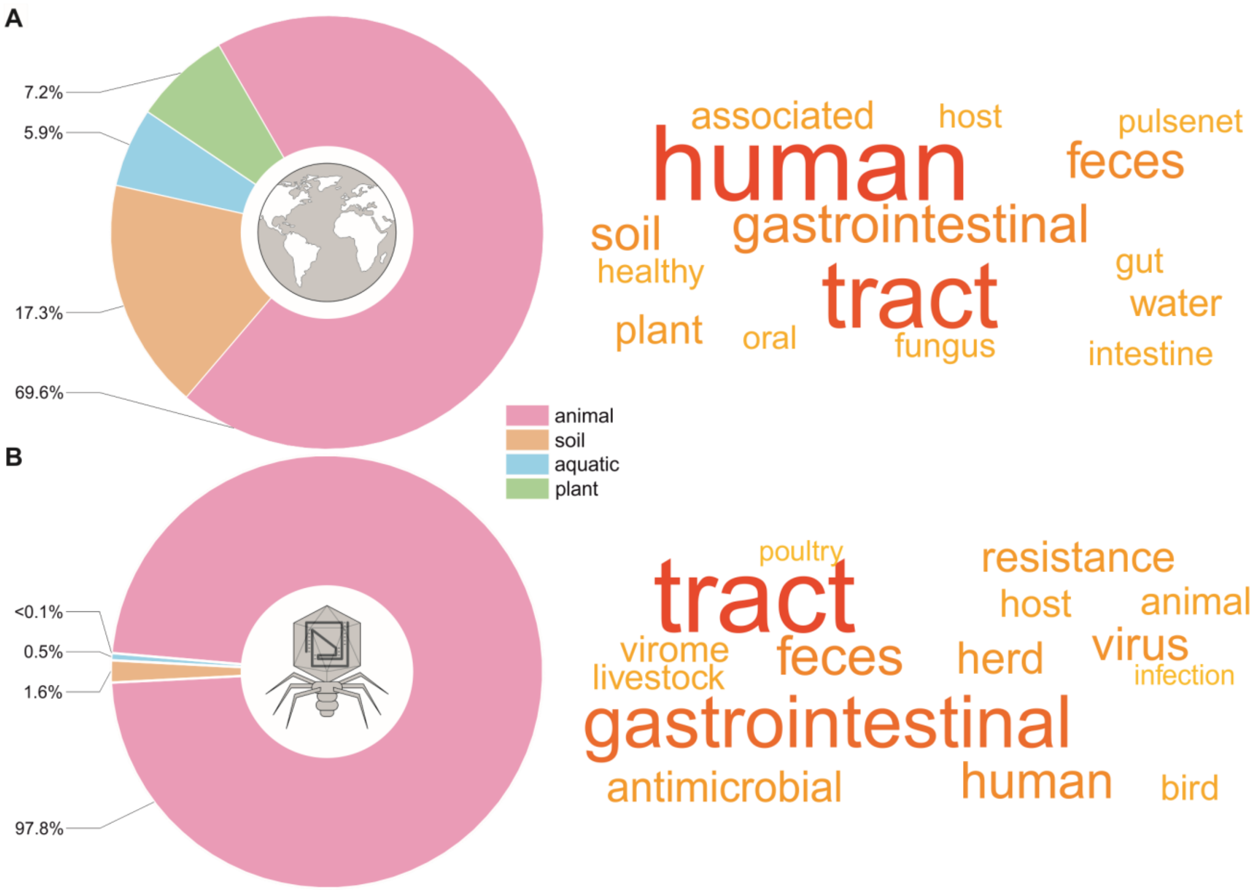


**Fig. S5. Animal gut is the main habitat for Θrzs.** Environments and keywords from annotated samples within the Microbe Atlas Project. The darkness and size of the words correlate with their occurrences (darkest, largest = most common). (**A**) Background distribution of keywords and environments of metagenomic samples within the Microbe Atlas Project: 10'000 samples were randomly selected, and the 15 most common keywords are shown. (**B**) The 15 most common keywords and environments from all samples in which tRNA-associated Θrzs are detected. Keywords related to viruses that were absent in the background distribution are now the top hits, whereas keywords such as "water", "plant" or "soil" are absent. The resulting word cloud point to viruses in the animal gut as the predominant Θrz environment.

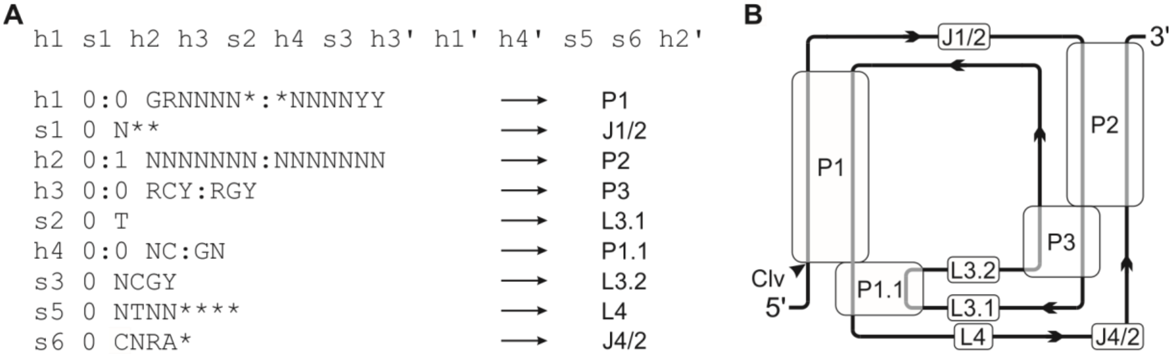


**Fig. S6. Structure of the refined Θrz descriptor file.** (**A**) Structure of the descriptor file of the final motif used in the RNArobo (*25*) motif search with corresponding domain names of the minimal drz motif. (**B**) Graphical illustration of minimal drz secondary structure and domain names for reference.

**Table S1. Summary of the investigated databases in the search for Θrzs.** Amount of detected Θrzs and tRNA-associated Θrzs in eight different bacteriophage databases. Representative genomes from proGenomes3 and eukaryotic genomes are also shown at the bottom.

| name of database | total no. of isolated Θrzs | total no. of tRNA-associated Θrzs | total no. of analyzed bp (Gbp) | origin |
|---|---|---|---|---|
| Tisza, 2021 (*27*) | 88 | 58 | 6.4 | human gut |
| Camarillo-Guerrero, 2021 (*48*) | 342 | 241 | 10.7 | human gut |
| Nayfach, 2021 (*50*) | 565 | 364 | 17.6 | human gut |
| Borges, 2022 (*22*) | 26 | 16 | 0.1 | animal gut |
| Li, 2022 (*51*) | 2 | 2 | 2.9 | human oral |
| Gregory, 2019 (*52*) | 7 | 0 | 7.2 | marine |
| Stano, 2016 (*49*) | 0 | 0 | 0.8 | all environments |
| Roux, 2021 (*53*) | 239 | 60 | 97.1 | all environments |
| proGenomes3* (*30*) | 12 | 0 | 310.6 | bacterial genomes |
| NCBI RefSeq* (*55*) | 0 | 0 | 7.8 | eukaryotic genomes |

*non-viral databases

**Table S2. Apparent $Mg^{2+}$-dependent cleavage rate constants ($k_{obs}$) of selected tRNA/Θrz pairs.** [$Mg^{2+}$] = Magnesium(II) concentration (mmol/L); $k_1$ = apparent cleavage rate constant ($min^{-1}$); rep1-3 = replicate 1-3; mean($k_1$) = mean value of $k_1$ ($min^{-1}$); σ = standard deviation of the mean ($min^{-1}$); $k_{max}$ = maximal apparent cleavage rate constant ($min^{-1}$); SE = standard error of the fit ($min^{-1}$); $K_d$ = dissociation constant (mM); n = Hill coefficient.

**$tRNA^{Leu}$0024_Θ0009**

| [$Mg^{2+}$] | $k_1$(rep1) | $k_1$(rep2) | $k_1$(rep3) | mean($k_1$) ± 3σ | $k_{max}$ ± SE | $K_d$ ± SE | n ± SE |
|---|---|---|---|---|---|---|---|
| 0.03 | 0.0028 | 0.0029 | 0.0024 | 0.0027 ± 0.0007 | 1.11 ± 0.27 | 1.51 ± 0.60 | 1.75 ± 0.21 |
| 0.1 | 0.010 | 0.006 | 0.008 | 0.008 ± 0.007 | | | |
| 1 | 0.48 | 0.52 | 0.49 | 0.50 ± 0.07 | | | |
| 10 | 0.94 | 1.02 | 1.09 | 1.02 ± 0.22 | | | |

**$tRNA^{Leu}$0114_Θ0016**

| [$Mg^{2+}$] | $k_1$(rep1) | $k_1$(rep2) | $k_1$(rep3) | mean($k_1$) ± 3σ | $k_{max}$ ± SE | $K_d$ ± SE | n ± SE |
|---|---|---|---|---|---|---|---|
| 0.03 | 0.0054 | 0.0063 | 0.0065 | 0.0061 ± 0.0019 | 2.32 ± 0.43 | 0.84 ± 0.27 | 1.88 ± 0.22 |
| 0.1 | 0.030 | 0.036 | 0.037 | 0.034 ± 0.011 | | | |
| 1 | 1.94 | 2.58 | 2.59 | 2.37 ± 1.13 | | | |
| 10 | 1.64 | 2.44 | 2.61 | 2.23 ± 1.55 | | | |

**$tRNA^{Sup}$0028_Θ0093**

| [$Mg^{2+}$] | $k_1$(rep1) | $k_1$(rep2) | $k_1$(rep3) | mean($k_1$) ± 3σ | $k_{max}$ ± SE | $K_d$ ± SE | n ± SE |
|---|---|---|---|---|---|---|---|
| 0.03 | 0.00011 | 0.00009 | 0.00008 | 0.00009 ± 0.00004 | 1.20 ± 0.22 | 1.53 ± 0.23 | 2.38 ± 0.08 |
| 0.1 | 0.0029 | 0.0036 | 0.0043 | 0.0036 ± 0.0020 | | | |
| 1 | 0.30 | 0.29 | 0.29 | 0.29 ± 0.02 | | | |
| 10 | 1.28 | 1.18 | 1.22 | 1.22 ± 0.15 | | | |

**$tRNA^{Val}$0025_Θ0046**

| [$Mg^{2+}$] | $k_1$(rep1) | $k_1$(rep2) | $k_1$(rep3) | mean($k_1$) ± 3σ | $k_{max}$ ± SE | $K_d$ ± SE | n ± SE |
|---|---|---|---|---|---|---|---|
| 0.03 | 0.00006 | 0.00007 | 0.00020 | 0.00011 ± 0.00023 | >10 | - | - |
| 0.1 | 0.0006 | 0.0006 | 0.0019 | 0.0010 ± 0.0023 | | | |
| 1 | 0.58 | 0.45 | 1.05 | 0.69 ± 0.94 | | | |
| 10 | >10 | >10 | >10 | >10 | | | |

**Table S3. Apparent pH-dependent cleavage rate constants ($k_{obs}$) of selected tRNA/Θrz pairs.** [$Mg^{2+}$] = Magnesium(II) concentration (mmol/L); $k_1$ = apparent cleavage rate constant ($min^{-1}$); rep1-3 = replicate 1-3; mean($k_1$) = mean value of $k_1$ ($min^{-1}$); σ = standard deviation of the mean ($min^{-1}$); $k_{max}$ = maximal apparent cleavage rate constant at 0.5 mM $Mg^{2+}$ ($min^{-1}$); SE = standard error of the fit ($min^{-1}$); p$K_{a1}$ = p$K_a$ of the hydrated $Mg^{2+}$ ion; p$K_{a2}$ = p$K_a$ of the catalytic cytosine residue.

| **$tRNA^{Leu}$0114_Θ0016** | | | | | | | |
|---|---|---|---|---|---|---|---|
| pH | $k_1$(rep1) | $k_1$(rep2) | $k_1$(rep3) | mean($k_1$) ± 3σ | $k_{max}$ ± SE | p$K_{a1}$ ± SE | p$K_{a2}$ ± SE |
| 5.06 | 0.0073 | 0.0076 | 0.0074 | 0.0074 ± 0.0005 | 0.88 ± 0.03 | 9.0 ± 0.1 | 6.3 ± 0.1 |
| 6.05 | 0.25 | 0.26 | 0.31 | 0.27 ± 0.10 | | | |
| 6.97 | 0.73 | 0.76 | 0.78 | 0.75 ± 0.07 | | | |
| 8.01 | 0.72 | 0.73 | 0.74 | 0.73 ± 0.04 | | | |
| 9.01 | 0.43 | 0.44 | 0.44 | 0.44 ± 0.03 | | | |
| **$tRNA^{Sup}$0028_Θ0093** | | | | | | | |
| pH | $k_1$(rep1) | $k_1$(rep2) | $k_1$(rep3) | mean($k_1$) ± 3σ | $k_{max}$ ± SE | p$K_{a1}$ ± SE | p$K_{a2}$ ± SE |
| 5.06 | 0.052 | 0.086 | 0.055 | 0.064 ± 0.056 | 0.75 ± 0.06 | 8.7 ± 0.2 | 5.8 ± 0.2 |
| 6.05 | 0.64 | 0.44 | 0.42 | 0.50 ± 0.37 | | | |
| 6.97 | 0.76 | 0.83 | 0.69 | 0.76 ± 0.21 | | | |
| 8.01 | 0.49 | 0.49 | 0.58 | 0.52 ± 0.15 | | | |
| 9.01 | 0.30 | 0.29 | 0.31 | 0.30 ± 0.04 | | | |
| **$tRNA^{Val}$0025_Θ0046** | | | | | | | |
| pH | $k_1$(rep1) | $k_1$(rep2) | $k_1$(rep3) | mean($k_1$) ± 3σ | $k_{max}$ ± SE | p$K_{a1}$ ± SE | p$K_{a2}$ ± SE |
| 5.06 | 0.0079 | 0.0068 | 0.0073 | 0.0073 ± 0.0017 | 0.084 ± 0.003 | 9.0 ± 0.1 | 6.2 ± 0.1 |
| 6.05 | 0.032 | 0.030 | 0.030 | 0.031 ± 0.003 | | | |
| 6.97 | 0.079 | 0.080 | 0.074 | 0.078 ± 0.009 | | | |
| 8.01 | 0.070 | 0.070 | 0.067 | 0.069 ± 0.005 | | | |
| 9.01 | 0.044 | 0.041 | 0.043 | 0.043 ± 0.005 | | | |

**Data S1. (separate file)**

Table containing the combined tRNA-associated Θrz sequences from annotated and metagenomic samples sorted by Θrz frequency in descending order. Contains tRNA-associated Θrzs, the respective tRNA sequences, and additional information.

**Data S2. (separate file)**

Table containing the combined isolated Θrz sequences from annotated and metagenomic samples sorted by Θrz frequency in descending order. Contains non-tRNA-associated Θrz sequences and additional information.

**Data S3. (separate file)**

Comma separated identifiers of raw reads from runs analyzed in the scope of the metagenomic search. All datasets are searchable and downloadable by the identifier (https://www.ncbi.nlm.nih.gov/sra).
